# Microbial Diversity and Community Shifts in a Petroleum Reservoir under Production: Effects of Water Breakthrough and Anthropogenic Alterations

**DOI:** 10.1101/2025.06.13.659474

**Authors:** Armando Alibrandi, Julia Plewka, Rolando di Primio, Alexander Bartholomäus, Aurèle Vuillemin, Alexander J. Probst, Jens Kallmeyer

## Abstract

Subsurface petroleum reservoirs host indigenous microorganisms that survive extreme conditions and long-term isolation. Microbial activity in these environments can contribute to adverse effects such as oil biodegradation and reservoir souring. Unlike the broader deep biosphere, oil reservoirs are frequently subjected to anthropogenic disturbances, particularly during production processes like water injection, which introduces external microbes and electron acceptors.

In this study, we investigated microbial diversity, community structure, and the impact of water breakthrough in the Edvard Grieg oil reservoir offshore Norway using 16S rRNA gene and metagenomic sequencing.

We found clear regional heterogeneity in community composition, with low diversity dominated by thermophilic, anaerobic, and halotolerant taxa. The southern region (wells A13, A17, A18, and A19) exhibited lower diversity, while well A07 showed a distinct microbial signature. The dominant genera included the strictly anaerobic bacterium *Thermoanaerobacter* and the hyperthermophilic archaeon *Thermococcus*. Water breakthrough triggered shifts in community structure, not due to widespread replacement by injected microbes, but via the increase of sulfate-reducing bacteria. Metagenomic data supported these observations and suggested rapid microbial transport between injection water and the water separator. These findings support the use of microbial signatures as cost-effective tools for monitoring oil reservoir processes and integrity.

## Introduction

Microbial communities are powerful agents of environmental transformation, capable of altering their surroundings through metabolic activity (Gupta et al., 2017). One of the most economically disruptive microbial processes in oil reservoirs is oil souring, the production of toxic and corrosive hydrogen sulphide (H₂S) by microbial sulphate reduction, leading to diminishing oil value (Gieg et al., 2011; Magot, 2005; Medina-Bellver et al., 2005). In addition to souring, microbial activity degrades the composition and properties of crude oil (Head et al., 2003). Biodegradation increases oil density, which is measured by the oil industry as API gravity (American Petroleum Institute), a key indicator of oil quality. Lower API values indicate higher density and lower quality.

A plethora of studies have examined the microbial composition of oil reservoirs (Cai et al., 2015; Lin et al., 2014; Ren et al., 2011; Takahata et al., 2000; Tang et al., 2012; Vigneron et al., 2017), yet most of them focused on production water samples from flooded reservoirs in which water-breakthrough had already occurred. In the oil industry, flooding is the process where fluid injections, mostly seawater for offshore reservoirs as well as formation water, are used to maintain pressure during the extraction process and increase recovery rates (Clemens et al., 2017). Formation water, naturally present in the reservoir, is produced along with crude oil and is separated from the oil onboard the oil production platform. Once the injected water reaches the production well, i.e. the so-called water breakthrough occurs, recovered oil becomes mingled with the injected fluids. Under such circumstances, microbial community analyses may reflect introduced taxa rather than indigenous reservoir populations. However, even in non-water flooded reservoirs, contamination from drilling, well operations, or faulty tubing potentially introduces foreign microorganisms, raising the question of whether the microorganisms driving biodegradation and souring are truly native or introduced through drilling or well operations (Magot et al., 2000; Wentzel et al., 2013; Youssef et al., 2009).

Water injection can transport non-native microorganisms into the reservoirs. Even small amounts of seawater can drastically alter microbial communities (Li et al., 2017; Vigneron et al., 2017) due to its significantly higher microbial abundance (10^3^ to 10^9^ cells/ml; (Bar-On et al., 2018; Jannasch and Jones, 1959; Kong et al., 2020, p.202) compared to the typical in-reservoir densities (∼10⁴ cells/ml, depending on temperature and salinity; (Magot, 2005). Additionally, water injections into an oil field induce a reduction of *in-situ* temperatures, thereby potentially fostering environmental conditions conducive to microbial proliferation within the reservoir (Vigneron et al., 2017). By adding sulphate, nitrogen, and phosphorus sources, seawater injection sets the stage for the growth of sulphate-reducing bacteria (SRB), with subsequent production of H_2_S (Vance and Thrasher, 2005).

Beyond their detrimental impact on the value of the oil, microbial communities also hold promise as biosignatures for understanding reservoir dynamics. Recent studies demonstrate that microbial composition can serve as a tracer of oil migration pathways and fluid provenance, often providing greater precision than traditional geochemical analyses (Zhang et al., 2020, 2021). As sequencing technologies become increasingly cost-effective, microbial profiling is emerging as a practical tool for oil field monitoring (DNA Sequencing Costs: Data, 2024).

To date, the upper-temperature limit for microbial life is currently documented at 122°C, as demonstrated for *Methanopyrus kandleri* (Takai et al., 2008). However, in oil reservoirs, biodegradation ceases above ∼80°C, as this process relies on microbial consortia where disruptions in metabolic linkages can halt the entire process (Head et al., 2014; Magot, 2005; Wilhelms et al., 2001). The exact cause of this temperature threshold remains uncertain, though hypotheses suggest that energy demands for cellular repair in nutrient-limited reservoir environments become prohibitive (Larter et al., 2003). Recently, a study reported that as microbial cells approach the upper-temperature limit of life in deep, hot subsurface sediments, cellular metabolic rates increase again to counter the thermal degradation of biomolecules (Beulig et al., 2022). Moreover, hydrocarbons impose solvent stress on cell membranes, a challenge that intensifies with rising reservoir temperatures (Pannekens et al., 2019). Combined with high salinity and the abundance of heavy metals, these extreme conditions select for specialized poly-extremophiles (Youssef et al., 2009), primarily from anaerobic and thermophilic clades of Firmicutes, Euryarchaeota, and Thermotogota (Table 1; Wentzel et al., 2013). While some members of these groups, such as *Thermococcus*, can thrive at temperatures exceeding 80°C (Zhao et al., 2015), the combination of multiple stressors in oil reservoirs appears to prevent microbial communities from actively degrading hydrocarbons under such conditions.

**Table 1:** Phylum and genera considered to be indigenous to hot oil reservoirs. (Head, 2017; Vigneron et al., 2017; Wentzel et al., 2013). New(er) nomenclature in parenthesis.

| Phylum | Genera |
| --- | --- |
| Thermotogota | *Petrotoga*, *Kosmotoga*, *Thermotoga*, *Geotoga*, *Oceanotoga*,<br>*Thermosipho* |
| Firmicutes<br>(Bacillota) | *Thermoanaerobacter*, *Geobacillus*, *Bacillus*, *Desulfotomaculum*,<br>*Caldanaerobacter*, *Mahella*, *Caminicella* |
| Deinococota | *Thermus* |
| Synergistetes<br>(Synergistota) | *Thermovirga*, *Anaerobaculum* |
| Deferribacteres(<br>Deferribacterota<br>) | *Deferribacter* |
| Euryarchaeota<br>(Methanobacter<br>iota) | *Methanoculleus, Methermicoccus, Methanothermobacter,*<br>*Methanococcus, Archaeoglobus, Thermococcus* |

Oil reservoir temperatures, dictated by sedimentary depth and geothermal gradients, range from ∼20°C to 150°C (Peters et al., 2007, p.200). With increasing burial depth, temperatures typically rise at about 3°C per 100 meters (Magot, 2005), and in deeper formations, they may exceed 130 to 150°C, with hydrocarbon accumulations predicted at depths of up to 13 km (Pang et al., 2022, p.20). Here, we define “hot reservoirs” as those conditions approaching the 80°C limit for biodegradation (Wilhelms et al., 2004) rather than the absolute upper temperature limits of microbial life.

Wentzel et al. (2013) compiled a list of microbial taxa that are most likely indigenous to reservoirs, while others are most likely introduced contaminants from drilling and oil production processes.

Given the pivotal role that microbial communities play in biogeochemical processes within petroleum reservoirs, comprehending their responses to anthropogenic interventions - such as water injection - is paramount in monitoring well evolution during production. Our study aims to elucidate the spatial distribution of microbial communities in Edvard Grieg, a high-temperature (∼78°C) oil reservoir located in the North Sea and assess the effects of water-breakthrough on the structure of the indigenous microbial community in the reservoir.

## Materials and Methods

### Study site and well sampling for oil and water

Oil and water samples were collected from the Edvard Grieg oil field, located on the Norwegian continental shelf approximately 200 km West of Stavanger, Norway. The reservoir is located at an average depth of 1900 m below the seafloor (mbsf), and *in-situ* temperatures range between 76°C and 78°C. It is geologically subdivided into three segments, i.e. Luno, Jorvik, and Tellus, which are composed of sandstones, conglomerates and in part also weathered granitic basement rocks. Lacustrine shales are non-reservoir sediments in the field (Fig. 1). Edvard Grieg was discovered in 2007 and has been in production since 2015.

**Figure 1:**
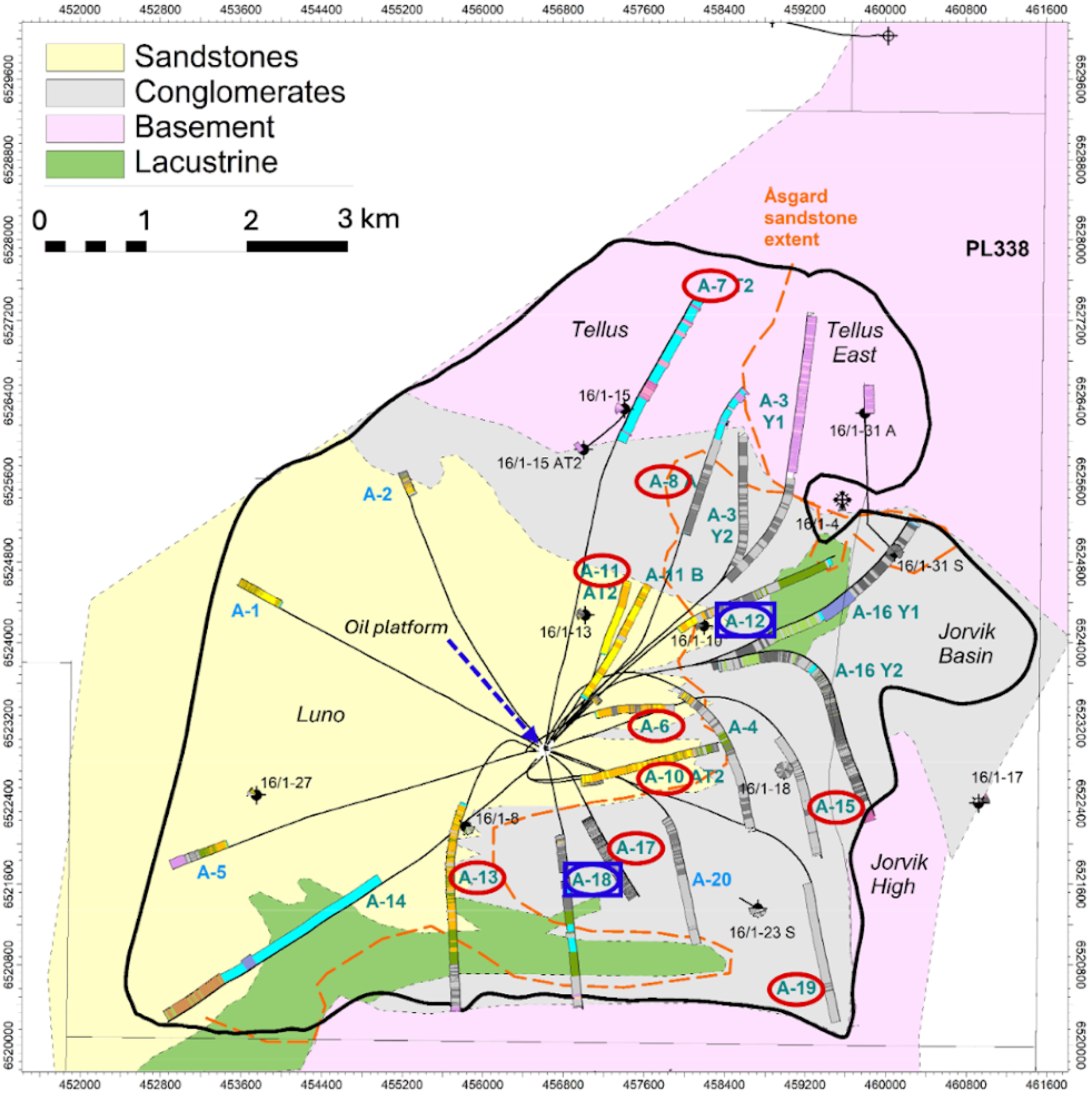
Map of the Edvard Grieg reservoir. The “A” series marks denote sampled well numbers. Blue squares represent sites sampled both before and after the water breakthrough, while red circles indicate sites sampled only before the water breakthrough. The black dot at the centre of the flow lines marks the location of the oil platform.

Produced fluids samples, i.e. crude oil that did not undergo water separation, were collected from eleven wells (A6, A7, A8, A10, A11, A12, A13, A15, A17, A18, A19). Each of these wells is directly connected to the Edvard Grieg platform (Fig. 1), where the crude oil mixed with suspended water is subjected to phase separation. On the day of sampling the oils from the different wells had a water cut (i.e. fraction of water suspended in the oil) ranging from 1% to 36% (Suppl. Table S9) and had API gravity values of 32 to 34. The high API values of Edvard Grieg correspond to a low density, which can be interpreted as the oils having a low level of biodegradation, which is supported by geochemical analyses.

During the first round of sampling, the reservoir had its pressure maintained by water injections but had not yet undergone a water breakthrough. Sixteen months after the first sampling and one month after the water breakthrough, we conducted a second round of sampling. Produced fluids were collected from wells A12 and A18, along with samples of separated oil, separated water, and injection water. The injection water consisted of reservoir formation water that was separated from the oil, mixed at variable proportions with oxygen-depleted seawater. During injection, deposition inhibitors and biocides were also added, and injection water temperature ranged from 40°C to 50°C.

Crude oil samples were collected from sampling valves located on the wellhead and directly transferred into sterile glass bottles. Bottles were filled without headspace to maintain anaerobic conditions. They were kept at room temperature during transport and stored at 4°C at GFZ Potsdam until analysis. For DNA extraction from crude oil, 25 ml of oil was transferred into 50 ml centrifuge tubes in an anaerobic glovebox to avoid sample exposure to oxygen. Further handling was carried out in a fume hood to avoid exposure to volatile hydrocarbons from the oils and various solvents used during the DNA extraction procedure (Alibrandi et al. 2023). Each experiment was carried out in triplicate.

### DNA extraction

DNA was extracted following the modified isooctane method described in Alibrandi et al. (2023). Briefly, samples were processed by mixing 25 ml of produced fluids with an equal volume of isooctane (2,2,4-trimethylpentane), followed by centrifugation at 5000 × g). After centrifugation, the supernatant was discarded, and the pellet was added to the bead tubes of the DNEasy PowerSoil Pro kit (Qiagen, Hilden, Germany) for DNA extraction. We added 40 µl of 10 % sodium dodecyl sulphate (SDS) to each bead tube and incubated the sample slurry at 65 °C for 10 minutes. DNA extraction was then processed according to the manufacturer’s instructions.

Production and injection water samples were filtered using Sterivex filters with a pore size of 0.22 µm (Merck KGaA, Darmstadt, Germany) to collect the suspended material. The filters were directly added to the bead tubes with 40 µl of 10 % SDS, heated at 65 °C for 10 minutes, and processed following the DNEasy PowerSoil Pro kit protocol. DNA concentrations in the final extracts were quantified using the Qubit 2.0 fluorometer with the dsDNA HS dying assay (Invitrogen, Carlsbad, USA).

### Amplification of 16S rRNA genes and amplicon sequencing

All samples were processed in duplicates. Bacterial and archaeal 16S rRNA gene fragments (V4 hypervariable region) were PCR amplified with the universal barcoded primer pair 515F (5′-GTG TGY CAG CMG CCG CGG TAA-3′) and 806R (5′-CCG GAC TAC NVG GGT WTC TAA T-3′). The final volume of each reaction mixture was 50 µl, consisting of 2 µl DNA template, 0.5 µl Taq DNA polymerase, 2 µl dNTP, 2 µl MgCl_2_, 5 µl 10 × polymerase buffer, 0.5 µl BSA, 2.5 µl of each primer, and 33 µl PCR grade water (EurX, Gdansk, Poland). PCR amplification was run at 95 °C for 5 min of initial denaturation, followed by 32 cycles of 30 sec at 95 °C (melting), 30 sec at 56 °C (annealing), and 1 min at 72 °C (elongation), with a final elongation of 7 min at 72 °C. PCR products were cleaned using AMPure magnetic beads (Beckman Coulter, Brea, USA), and barcoded samples were normalized to 20 ng of DNA and pooled. Amplicon sequencing was performed on an Illumina MiSeq platform using 2 × 300 base pair (bp) reads at Eurofins Genomics (Ebersberg, Germany).

### 16S rRNA data sequencing data treatment and statistical analysis

Read demultiplexing was performed using Cutadapt (v. 3.5; Martin, 2011) with the following parameters: -e 0.2 -q 15,15 -m 150 --discard-untrimmed. The amplicon sequence variants (ASVs) were generated using trimmed reads and the DADA2 package (v. 1.20; Callahan et al., 2016) with R v. 4.1, applying the pooled approach with the following parameters: truncLen = c(220,180), maxN = 0, rm.phix = TRUE, minLen = 160. Taxonomic assignment was done using DADA2 against the SILVA 16S rRNA SSU database release 138 (Quast et al., 2012). ASVs representing chloroplasts, mitochondria and singletons were removed. The partial 16S rRNA gene sequences were aligned using SINA Online (v.1.2.11 Pruesse et al., 2007) and inserted into a Maximum Likelihood RAxML phylogenetic tree on ARB (Ludwig et al., 2004). ASVs attributed to known bacterial extremophiles and SRB (92 ASVs) and archaea (99 ASVs) were selected and plotted into two separate phylogenetic trees, using the maximum parsimony algorithm with the bacterial and archaeal filters, and selecting the best tree among 100 replicates.

Statistical analyses of alpha and beta diversity were conducted using the Vegan community ecology package in R v. 4.3.1 and the software PAST 4.14 (Hammer et al., 2001). The dataset was rarefied to 2000 reads and included 3402 out of the total 3406 ASVs. Downstream statistical analyses were performed with the same rarefied dataset. For a general assessment of the microbial composition, we applied two key metrics: the relative read abundance of ASVs and α-diversity, including the observed richness and Shannon index. To examine regional differences within the reservoir, we looked at the beta-diversity of the microbial composition and conducted a Principal Coordinates Analysis (PCoA) with the Bray-Curtis similarity index, as well as a non-parametric Permutational Multivariate Analysis of Variance (PERMANOVA). The PCoA enabled visualization of the dissimilarity between microbial communities, providing insights into clustering patterns and trends across different regions. PERMANOVA was used to determine the statistical significance of the observed differences.

To investigate microbial community dynamics within the reservoir before and after injection, we combined PCoA and PERMANOVA approaches, as described above, with the Indicator Value (IndVal) Analysis to assess overall shifts in microbial composition and identify which specific microbial taxa can be significantly associated with the pre-injection, post-injection, water separator, oil separator, and water injection samples. This method considers both relative species abundance and occurrence frequency to determine the specificity and fidelity of a given group. For the calculation, we only used ASVs that had a cumulative count of more than 100, resulting in 29 selective ASVs.

To assess whether the water cut of the oils influenced the microbial composition, we ran a PERMANOVA test to see if the microbial composition varies with the water cut and ran a Mantel test to measure the correlation between the microbial structure and the water cut. Additionally, we performed non-metric multidimensional scaling (NMDS) based on Bray-Curtis dissimilarity to visualize patterns in microbial community composition across samples. The stress value was reported to indicate the goodness of fit of the ordination (<0.2 indicates a good fit). To test whether specific ASVs are influenced by the water cut a generalized additive model (GAMs) was applied (v. 1.9-3; Wood, 2017) using the R package mgcv. Significance was obtained by p-value < 0.05 corrected by the FDR method.

### Metagenomic sequencing, de novo assembly, and gene annotation

DNA extracts were sent to CeGaT GmbH (Tübingen, Germany) for metagenomic sequencing. Libraries were prepared using the Nextera XT DNA Library Preparation kit (Illumina), and sequencing was performed on a NovaSeq 6000 Illumina platform at CeGaT, aiming for 50 million read pairs (2 × 150 bps). Functional annotations of predicted Open Reading Frames (ORFs) were performed using the software package DIAMOND protein aligner (v. 0.9.24; Buchfink et al., 2015). Proteins were annotated with eggNOG (v. 2.1.12 and eggNOG DB v. 5.0.2; Cantalapiedra et al., 2021; Huerta-Cepas et al., 2019). We performed a quantitative functional annotation, focusing on open reading frames (ORFs) encoding proteins associated with sulphate, sulphite, and polysulphide reduction; nitrate and nitrite reduction; hydrocarbon degradation; methanogenesis; heat shock response; salt stress; biofilm formation; and microaerophilic respiration (Suppl. Tables S7). We ran a PERMANOVA analysis to assess whether the differences between the ORFs of interests associated with the sample groups were significant.

### Principal coordinate analysis based on gene sequence coverage

BBDuk (v. 37.09; Brian Bushnell, 2014) removed Illumina artefacts and adapters from the shotgun metagenomic raw reads. We trimmed the reads based on the quality scores with Sickle (v. 1.33; Joshi and Fass, 2011) and deduplicated them with BBMap (v. 37.09; Brian Bushnell, 2014). We assembled the quality-controlled reads with metaSPAdes (v.3.15.5; Nurk et al., 2017). Afterwards, all scaffolds were filtered with at least 1000 base pairs using pullseq (v. 1.0.2; Brian Thomas, Nilesh Patra, Tony Mancill, 2015), which were then used for gene prediction via prodigal (Hyatt et al., 2010). The nucleotide sequences of predicted genes from all metagenomes were clustered with MMSeqs2 (v. 15.6f452; (Steinegger and Söding, 2017) in cluster-mode 2, coverage-mode 1, minimum breadth of 95% and minimum sequence identity of 95%. Quality-controlled reads were mapped against the representative sequences of the resulting clusters with bowtie2 (v. 2.3.5.1; Langmead and Salzberg, 2012) in sensitive mode. Sequences were counted as present in samples if the minimum coverage breadth was greater than 95% and coverage greater than 5. The coverage values were normalized based on base pair counts of the sequenced forward reads. Only gene sequence clusters present in more than one metagenomic sample were used for the PCoA calculation to avoid the influence of undersampling. The PCoA was visualized with the R (R Core Team, 2020; RStudio Team, 2020) package ggplot (Hadley Wickham, 2016). The upSet plot was created with the package ComplexUpset (Michał Krassowski et al., 2022).

### Prokaryotic community composition based on extended rpS3 gene sequences

The ribosomal protein S3 (*rpS3*) marker gene was used to estimate the prokaryotic community composition (Sharon et al., 2015). Marker genes were identified with species-specific Hidden Markov Models (HMMs) and by comparing the amino acid sequences of predicted genes with diamond blastp (Buchfink et al., 2015) against the UniRef100 database (downloaded on 23.06.2021; The UniProt Consortium, 2019) with an e-value cut-off of 1e-5. The *rpS3* gene nucleotide sequences with 1000 bp flanking regions were extracted for all samples and clustered with MMSeqs2 (v. 15.6f452; Steinegger and Söding, 2017) in cluster-mode 2, coverage-mode 1, minimum breadth of 95% and minimum sequence identity of 95%. *RpS3* gene sequences were taxonomically annotated by comparing them with rpS3 sequences extracted from the GTDB (GTDB v. 220; Parks et al., 2022) with usearch -ublast (v. 10.0.240_i86linux64; Edgar, 2010). Sequences that could not be annotated via this approach, but that were binned, were assigned the MAG taxonomy (by finding the bin that carried the scaffold with the rpS3 gene sequence; for MAG construction, please see below). Quality-controlled reads of all samples were mapped against the representatives with bowtie2 (v. 2.3.5.1; Langmead and Salzberg, 2012) in sensitive mode. Reads mapping with more than ten per cent mismatches were excluded. The mean coverage depth of extended *rpS3* sequences was calculated for all sequences with a coverage breadth greater than 95%. Coverage was normalized by the base pair count of the forward reads. The data was visualized in R (R Core Team, 2020) with the Rstudio interface (RStudio Team, 2020). DNA extracts were sent to CeGaT GmbH (Tübingen, Germany) for metagenomic sequencing. Libraries were prepared using the Nextera XT DNA Library Preparation kit (Illumina), and sequencing was performed on a NovaSeq 6000 Illumina platform at CeGaT aiming for 50 million read pairs (2 × 150 bps). Due to low biomass in the samples from all wells, DNA concentrations were only sufficient to achieve library preparation and metagenomic sequencing on 6 samples, i.e. well A18 before injection, well A12 and A18 after injection, production water, and injection water.

### Reconstruction of metagenome-assembled genomes

Scaffolds with a minimum length of 1000 bp were binned into MAGs using ABAWACA (v. 1.0.0; Brown et al., 2015) and MaxBin2 (v. 2.2.7; Wu et al., 2016) with default parameters. The bins were aggregated with DASTool (v. 1.1.6; Sieber et al., 2018), and the resulting selection was manually curated in uBin (v. 0.9.14; Bornemann et al., 2020). Completeness and contamination of curated MAGs were calculated with CheckM2 (v. 1.0.1; Chklovski et al., 2023) and used GTDB-tk (v. 2.4.0; Chaumeil et al., 2022) with the Genome Taxonomy Database (v. 220; Parks et al., 2022) to assign taxonomy. The taxonomic annotation was used to build a de novo tree of the MAGs with GTDB-tk (Chaumeil et al., 2022) de_novo_wf workflow, converted into an iTOLs usable format with GTDB-tk’s convert_to_itol workflow and visualized with iTOLs (v. 6; Letunic and Bork, 2024).

ANI analysis was performed using FastANI (Jain et al., 2018) to compute the pairwise average nucleotide identity among the *Thermoanaerobacter* genomes. This tool identifies orthologous genomic fragments via bidirectional mappings and calculates the percentage of nucleotide similarity. Following established criteria, genomes with an ANI greater than 95% were considered to belong to the same species. Despite the fragmented nature of some assemblies, which may affect fragment ordering, the ANI values provide a robust measure of overall genomic similarity.

### Prediction of putative viral scaffolds and strain clustering

Viral strains are highly specific to their microbial hosts. To assess their potential as source tracers, we collected 31 putative viral strains from the samples and compared their presence/absence using a software suite (suppl. methods).

## Results

### Microbial diversity and spatial variation

Overall, the reservoir showed low taxonomic diversity, dominated by thermophilic and anaerobic taxa. The southern section of the Luno segment, represented by wells A13, A17, A18, and A19, showed lower microbial diversity compared to the rest of the reservoir. The most dominant taxa across all reservoir samples were attributed to the genera *Thermoanerobacter*, *Thermococcus,* and *Halomonas* (Fig. 2), accounting for most of the 16S rRNA gene abundance. Most ASVs were assigned to the genus level; the few that could not be assigned were removed from the plot.

**Figure 2:**
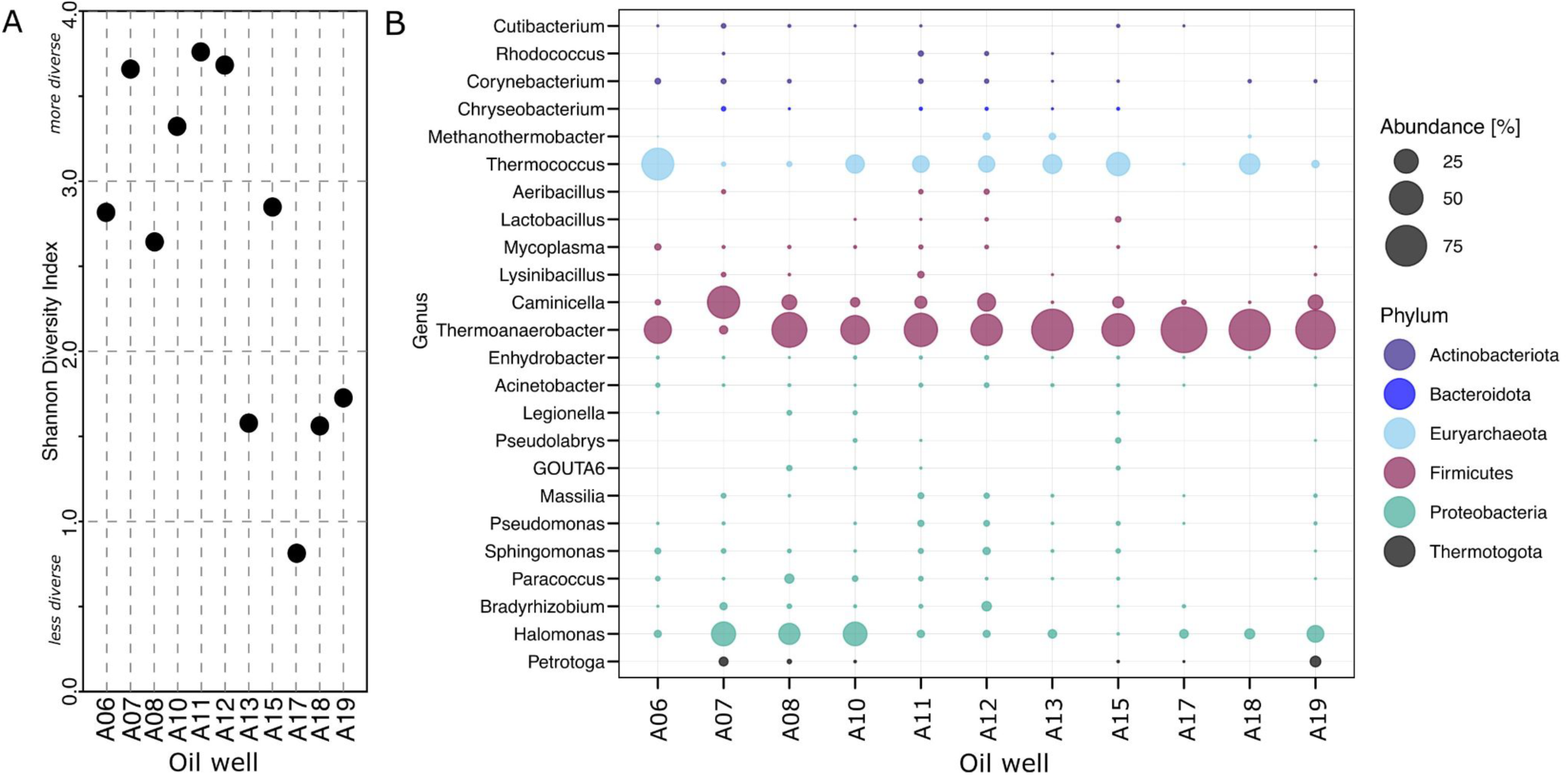
Abundance and diversity of the Edvard Grieg Reservoir. (A) Visualisation of the Shannon index. (B) Bubble plot shows 94% of the most abundant taxa. Each bubble is the average of the data points (n=3).

**Figure 3:**
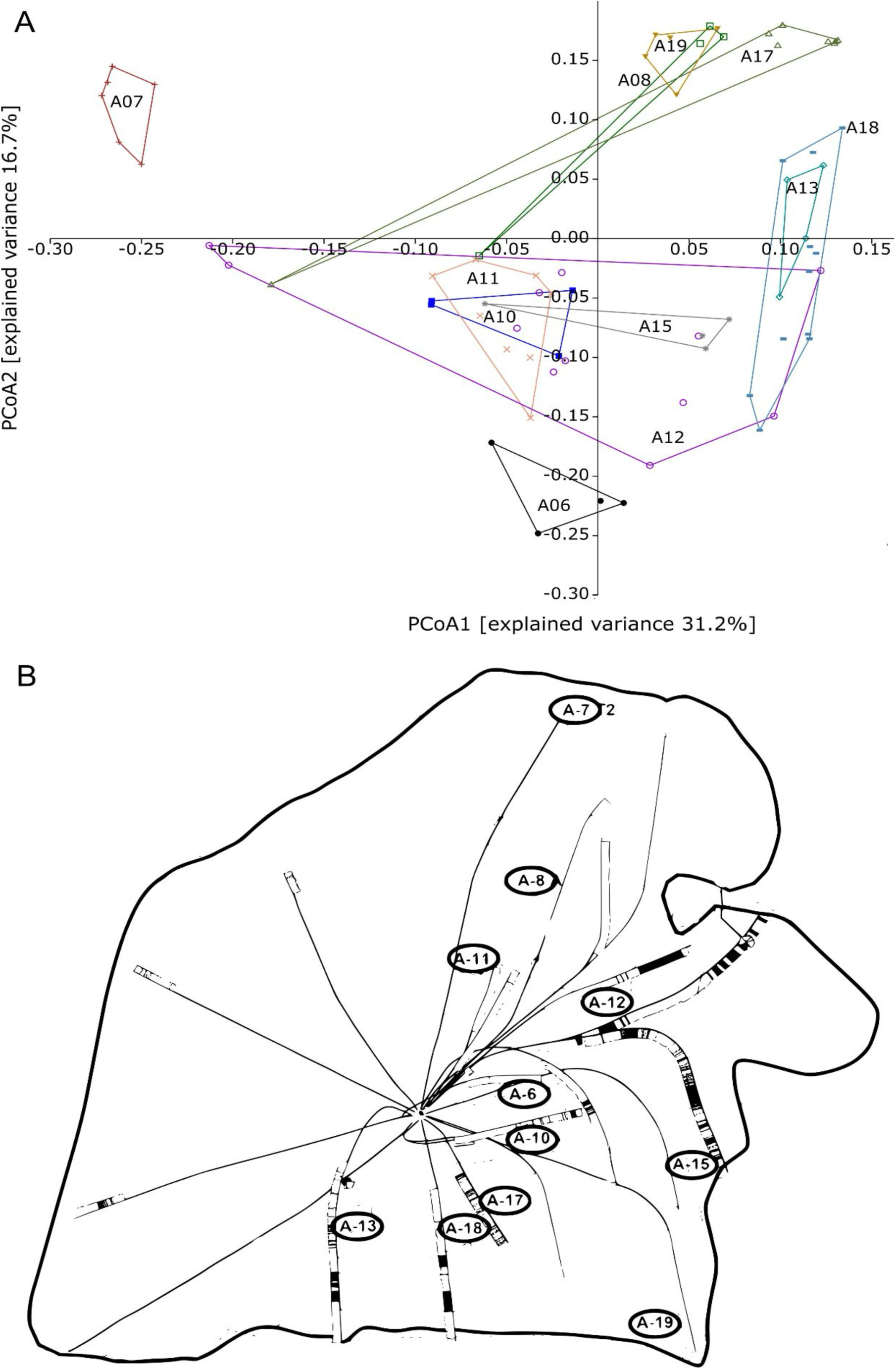
(A) Principal Coordinates Analysis (PCoA) plot of microbial communities from the Edvard Grieg reservoir, based on Bray-Curtis dissimilarity. Different symbols and colours denote the wellheads from which samples were collected. Axis 1 explains 31.2% of the variance, and Axis 2 explains 16.7%. (B) Simplified reservoir map illustrating the geographical layout of the wellheads, facilitating comparison with microbial community distribution.

### Impact of Water Breakthrough

Water breakthrough has a discernible impact on the microbial community composition within the oil reservoir. The PCoA plot (Fig. 4), with Coordinate 1 and Coordinate 2 showing 48.3% and 18.7% of the explained variance, respectively, demonstrates that the microbial communities before and after water-breakthrough exhibited statistically significant differences (p < 0.05; Suppl. Table S2). Before injection, oil samples from wells A12 and A18 exhibited a slightly higher degree of separation in the PCoA plot (Fig. 4) with a dissimilarity of 0.03. After water injection, the microbial communities in the oil samples from these two wells tended to shift towards uniformity as the dissimilarity index slightly decreased to 0.04. We infer from this observation a potential homogenising effect of water injection on the microbial community compositions. After the water breakthrough, the microbial community structure in the oil samples from the central well A18 (Fig. 1) shows higher similarity to those from the water separator, suggesting that major modifications in indigenous microbial communities originate mostly from microbes present in the platform, not from the injected water.

**Figure 4:**
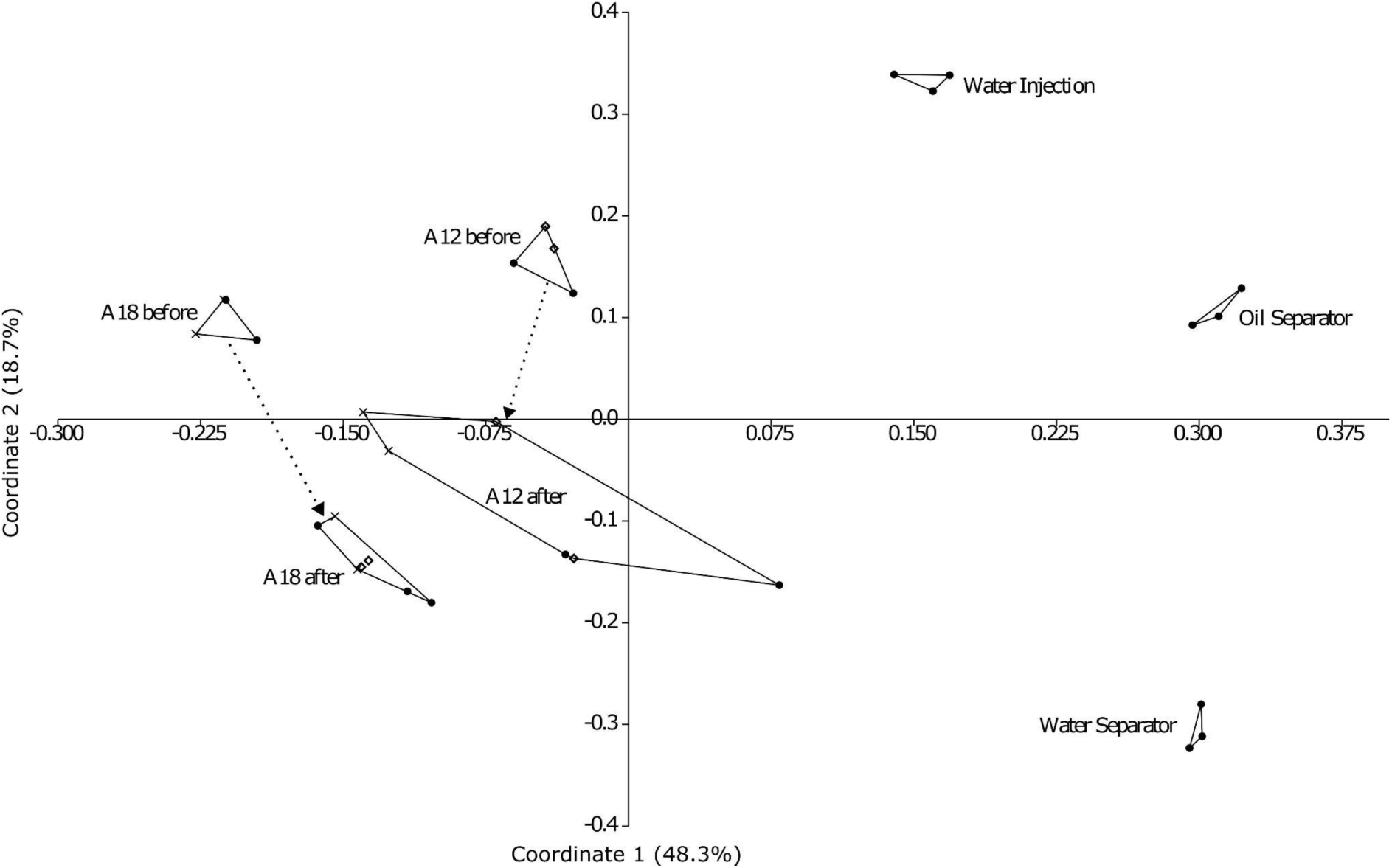
The effect of the fluid breakthrough on the microbial community. PCoA plot with Bray-Curtis similarity index of the two reference wells before and after injection, water injection and water separator. The different symbols correspond to the biological duplicates. Axis 1 48.3%, Axis 2 18.7% of explained variance.

Notably, the injection water, oil separator, and water separator samples exhibit no differences (p 0.09, 0.10; Suppl. Table S2). By contrast, the pre- and post-injection oils originating from A12 and A18 wells display significant differences (p 0.004). This suggests that the reservoir’s microbial communities underwent detectable changes between the two sampling time points

The four sulphate-reducer genera *Desulfofundulus, Archaeoglobus, Syntrophotalea* and *Desulfovibrio* were found exclusively in the production water, injection water and oil samples after injection (Fig. 5), suggesting that these taxa are not native to the reservoir, but were introduced through the injected water. By contrast, *Thermoanaerobacter, Caminicella, Methanothermobacter* and *Thermococcus* were consistently predominant across all samples, emphasising their tolerance to anaerobic hot conditions within the reservoir. These taxa were nevertheless also present in the injection water, suggesting that they may have been introduced during the injection process via the formation water mixed with seawater.

**Figure 5:**
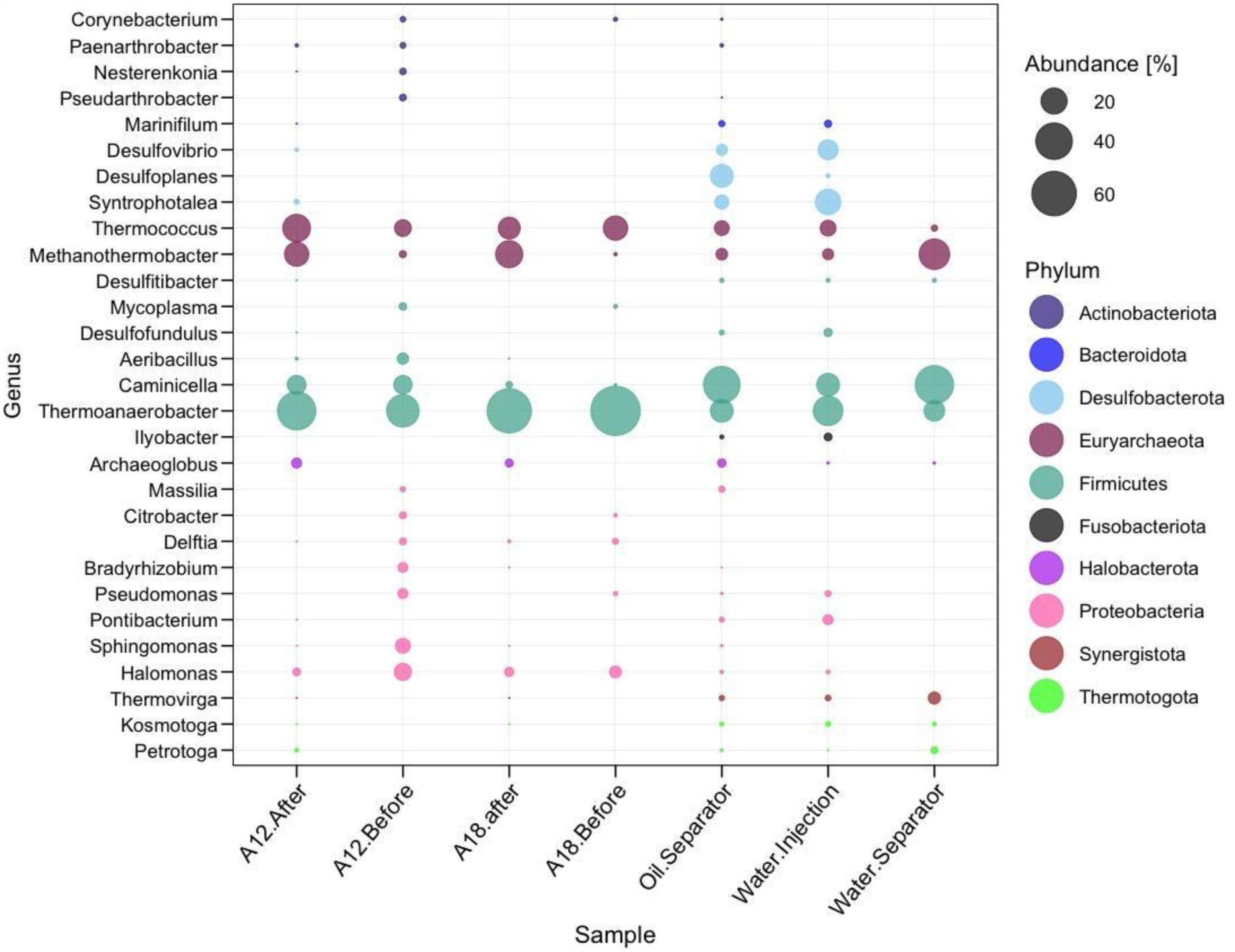
Bubble plot showing the relative abundance of 96% of the different genera present in the oil fluids before water injection, after injection, and, in the injection water (n=3).

The IndVal analysis (Fig. 6) identifies taxa with significant associations between specific samples. Water injection samples are significantly associated with multiple sulphate-reducing bacteria, such as *Desulfofundulus, Desulfovibrio* and *Synthrophotalea,* which likely indicates that these sulphate reducers were introduced during the water injection process. Taxa that likely represent contaminants from the oil processing include the genus *Dasulfoplanes* from the oil separator, and the genera *Petrotoga*, *Thermovirga* and a taxon from the order *Desulfomaculales* from the water separator (Fig. 5).

**Figure 6:**
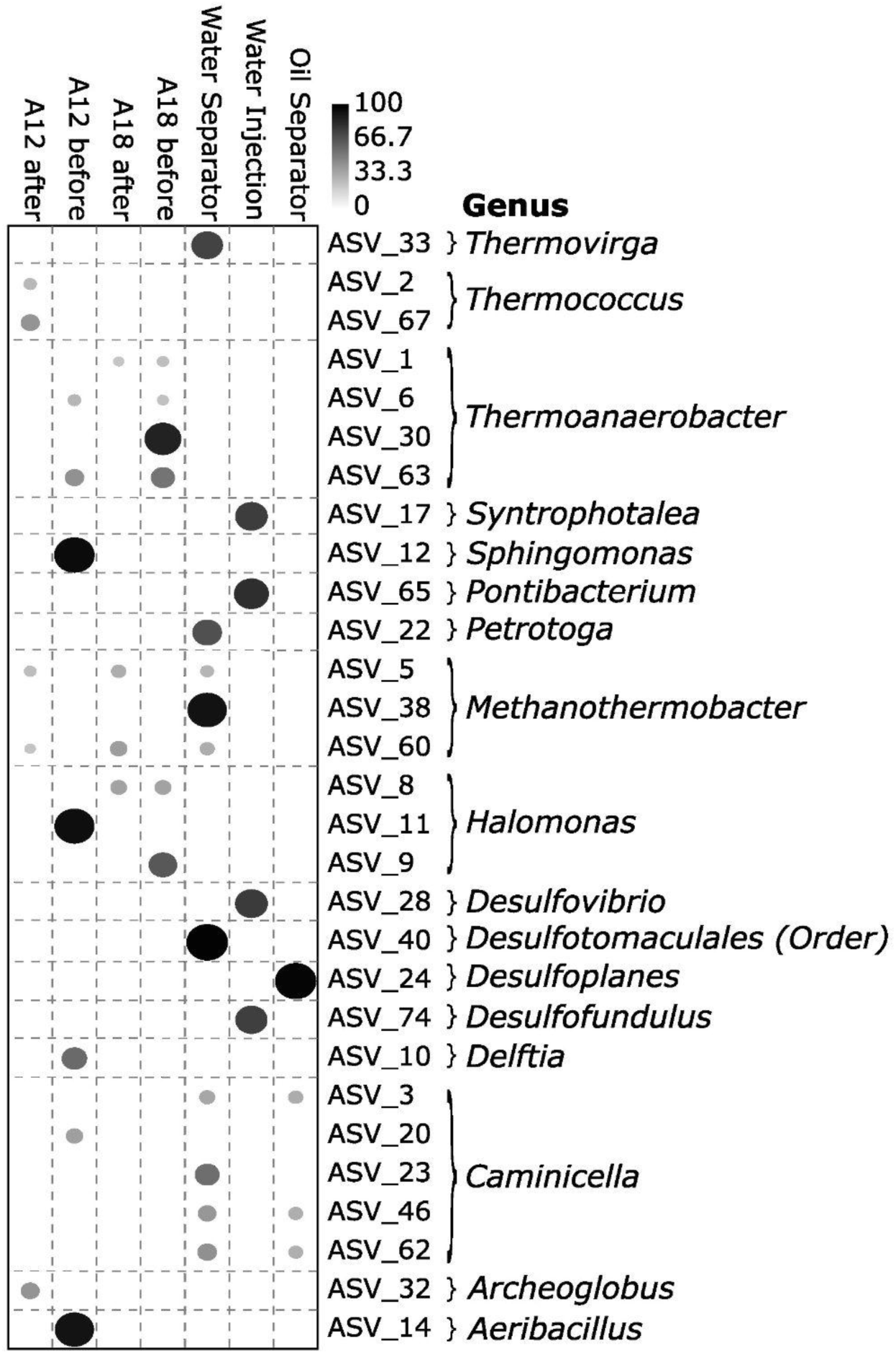
Indicator Species Analysis (IndVal) with a significance threshold of p<0.05. The dataset was rarified to 2000 reads, and only taxa with a cumulative count of more than 100 reads were selected for analysis and considered particularly indicative of specific samples. Higher IndVal percentages indicate stronger associations and closer links of selected taxa to specific sample origins.

The metagenomic analysis revealed a microbial community composition largely consistent with the 16S rRNA data. Community structure varied across samples, with distinct compositions observed before and after water injection, as well as in the water separator and water injection samples (Fig. 7). Across all samples, 248,889 genes were predicted, clustering into 177,477 species-level groups (Coelho et al., 2022), of which 20,769 were present in more than one sample. Notably, 28% of these shared clusters were detected in all samples (Suppl. Fig. 2).

**Figure 7:**
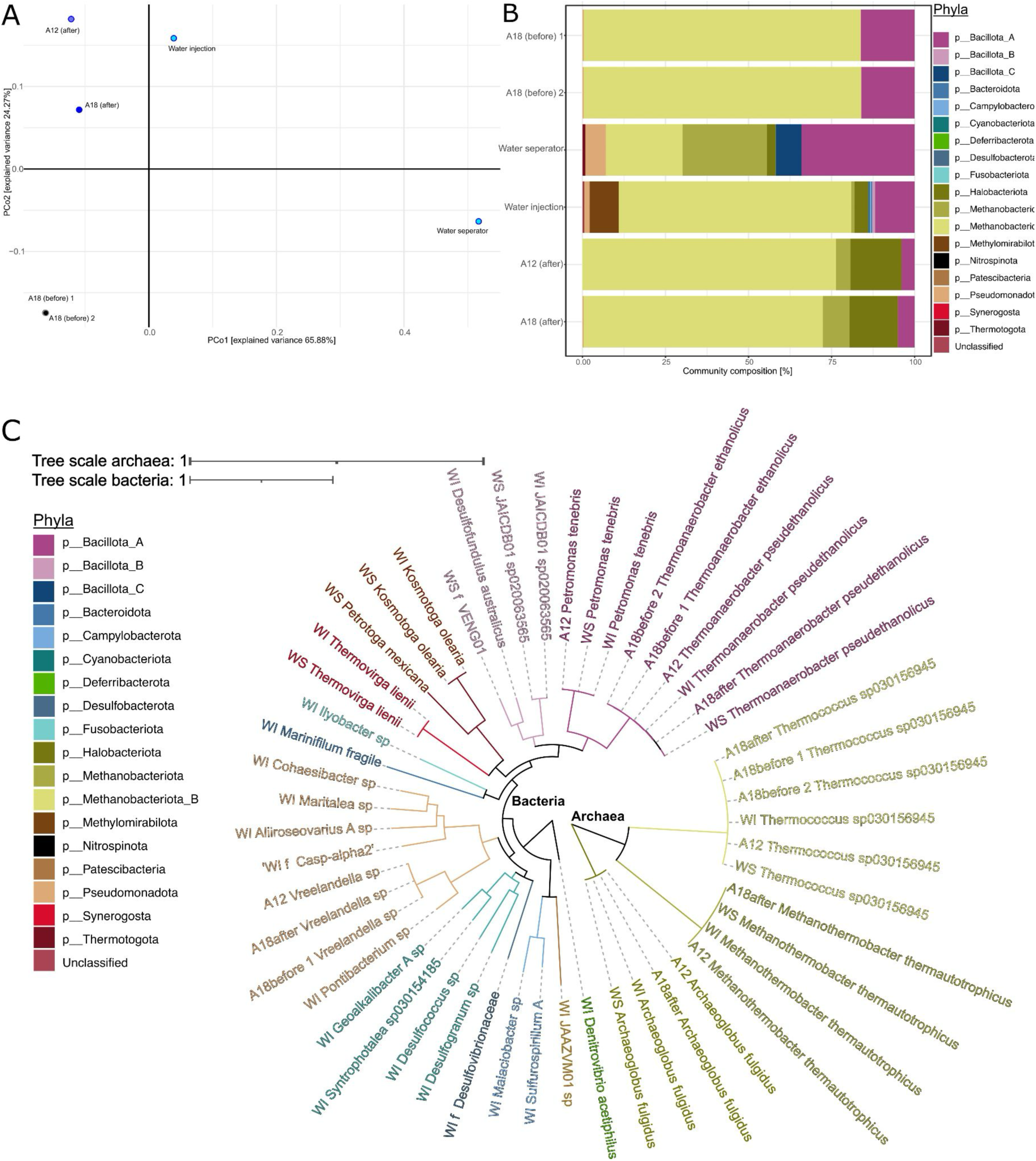
Metagenome and metagenome-assembled genome (MAG) characterisation in terms of functional and taxonomic diversity and read abundances. (A) 1 PCoA based on the abundance of 20,769 gene clusters. Oil samples taken before water was injected into the reservoir are coloured black, oil samples taken after water injection are coloured dark blue, and the samples of the water injection and water separator fluid are coloured light blue. (B) Community composition of prokaryotic organisms in the six samples based on coverage of extended rpS3 gene sequences. The proportion of phyla is shown by different colours. (C) Phylogenetic tree of metagenomic assembled genomes (MAGs). Nodes are coloured according to the phyla of the MAG. The acronym next to the taxon corresponds to where it was found: Water injection (WI), water separator (WS) and the wells (A12; A18).

Oil samples collected before water injection exhibited lower intra-group dissimilarity compared to those collected afterwards. Following water injection, the microbial community shifted, with samples clustering closer to the injection water sample. The water separator sample formed a distinct cluster, emphasizing its unique microbial composition. Although the dataset is statistically underpowered, these observations suggest that water injection has a strong impact on microbial community structure.

MAG quality assessment (Suppl. Fig. S3) showed that 27 out of 51 metagenome-assembled genomes (MAGs) met the high-quality threshold (≥95% completeness, <5% contamination). The majority of high-quality MAGs were reconstructed from the water injection sample (25), followed by the water separator (10). Key taxa, including Methanobacter_B, Bacillota_A, and Pseudomonadota, were retrieved from all sample groups. The relative abundance of MAGs (Fig. 7B) indicates substantial variation in microbial community composition between different sampling points. Before water injection, oil samples were dominated by *Thermococcus* (Methanobacteriota_B), with relative abundances reaching 69– 83%. After water injection, the proportion of *Thermococcus* decreased in the water separator sample (23%), reflecting a shift in community structure. Several phyla, including Bacillota_C, Methylomirabilota, and Desulfobacterota, were present in the water injection and water separator samples but were absent in the oil samples.

The genera *Archaeoglobus* (Halobacteriota) and *Methanothermobacter* (Methanobacteriota) were absent in the oil samples before injection but appeared in water and post-injection oil samples. *Methanothermobacter,* however, was found in the amplicon data prior to injection and this genus likely originates from the reservoir despite being associated with the water injection within the IndVal analysis. Taxa from the phyla Pseudomonadota and Bacillota_A were present across all samples but contributed differently to the overall community composition. Notably, the relative abundance of Bacillota_A, particularly RUG420 (Suppl. Fig. S2), declined 3.6-fold after water injection, suggesting a selective effect of the process.

Phylogenetic analysis (Fig. 7C) illustrates the taxonomic diversity of the microbial communities, including representatives from Euryarchaeota (Methanobacteriota), Desulfobacterota, Firmicutes (Bacillota), and Thermotogota. The presence of Desulfobacterota in water injection and water separator samples suggests a shift towards sulphate-reducing communities following water injection.

Metagenomic analysis from the viral strains revealed that well 18 had eleven viral strains in common with water samples after the water breakthrough compared to two viral strain clusters before, indicating a massive effect on the viral community after water injection (Suppl. Fig. S4). The sample prior to the water breakthrough from well A12 shares no viral strain clusters with water samples, but the presence of two viral strains from well A18 after the water injection is probably due to the use of formation water in the injection.

### The potential of functional genes

The ORFs selected from the metagenomes (Suppl. Table S7) revealed distinct patterns in functional gene abundance across samples taken before and after water breakthrough, highlighting shifts in microbial metabolic potential and stress responses associated with environmental changes (Fig. 8). Methanogenesis-related ORFs were undetectable in samples collected before water breakthrough but became evident in the water separator, injection water and post-water breakthrough oil samples. ORFs associated with hydrocarbon degradation were scarce and found mostly in post-water breakthrough samples.

**Figure 8.**
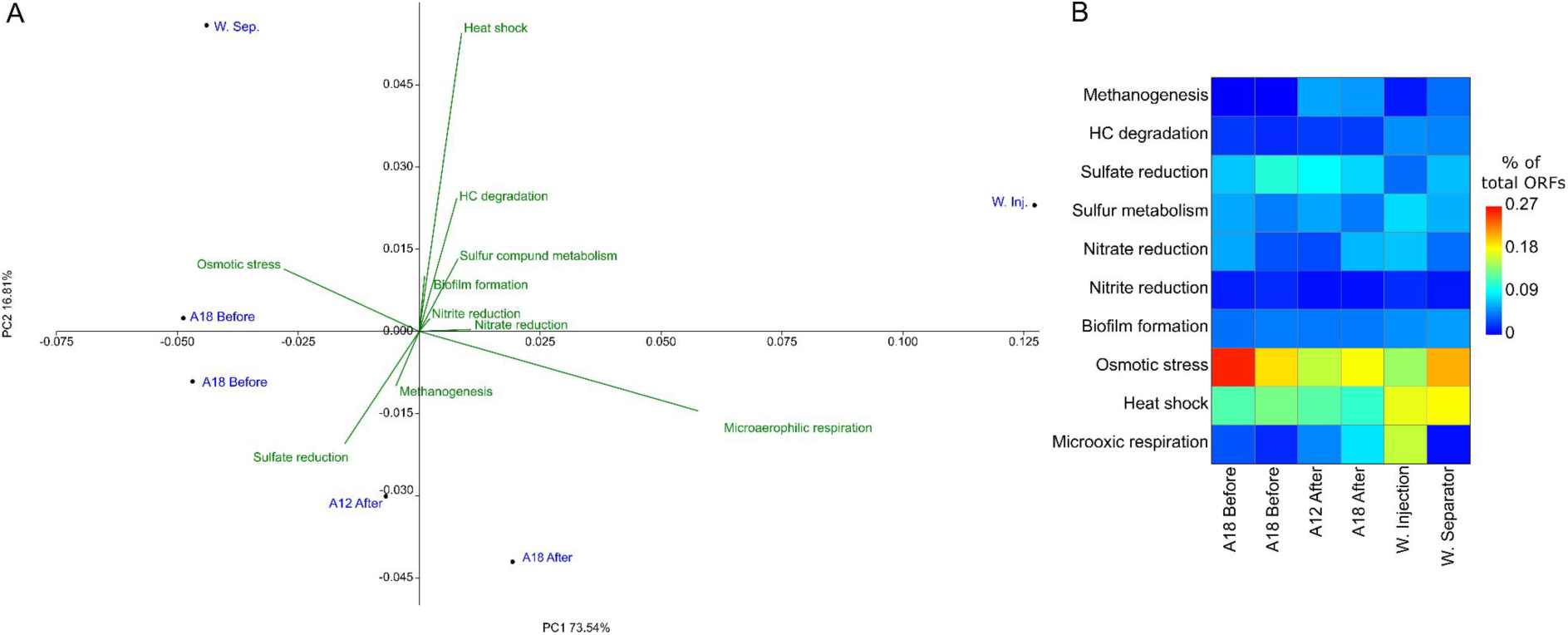
Distribution and abundance of functional genes relevant to oil reservoirs. (A) PCA biplot of the ORFs of interest, with green arrows indicating the direction of samples where the functional genes were found. (B) Heatmap showing the percentage of gene hits for the ORFs of interest relative to the total number of ORFs in each sample.

Sulphate reduction-related ORFs were abundant in all samples but scarce in the water injection sample. Nitrate and nitrite reduction were absent in the pre-breakthrough samples but present afterwards, with a particular abundance of ORFs in the injection itself. ORFs associated with osmotic stress were consistently detected across all samples, with the greatest abundance in one of the samples before the water breakthrough. Microaerophilic oxygen respiration-related ORFs were scarce before the water breakthrough and present in all remaining samples, with higher abundance in the water injection sample.

PERMANOVA results indicate that there is a statistically significant difference in ORF compositions between sample groups (p 0.0245), but the pairwise distance values show no significance based on ORFs composition.

## Discussion

### Regional differences and microbial composition

To the best of our knowledge, this study is the first to sample microbial communities directly from produced fluids, rather than from separated production water, across multiple locations within the same hot oil reservoir. The reservoir exhibited localised differences in microbial community composition, underscoring a crucial point: oil reservoir microbes could potentially serve as indicators for understanding oil provenance and migration pathways in the subsurface, provided that temperatures remain within suitable ranges. This approach, as suggested by Zhang et al. (Zhang et al., 2020) and supported by our data, presents a promising, low-cost complement to traditional geological and geophysical methods, enhancing our ability to monitor and understand reservoir dynamics in oil fields.

We assumed that, given the stringent DNA extraction protocol (Alibrandi et al., 2023), at least part of the DNA from endospores was also extracted and thereby sequenced with the total environmental DNA. Harsh reservoir conditions result in low biomass and, consequently, low DNA yields, estimated at 10²–10³ cells per mL based on per-cell DNA weight (Alibrandi et al., 2023). Direct cell counting is impractical due to low contrast between cells and oil, coupled with oil interference in fluorescence microscopy (Kallmeyer, 2011; Lloyd et al., 2013), which hampers reliable detection.

Microbial diversity across all wells of the reservoir was generally low. The oil samples A13, A17, A18, and A19 from the southern segment exhibit an even more pronounced reduced diversity. While the cause of this pattern remains unclear, the lack of correlation between water cut data and diversity indices suggests that local lithology, rather than water content, may be responsible. The microbial taxa identified in our dataset align with previous findings from oil reservoirs. Genera commonly reported in earlier studies and observed in our study include *Corynebacterium, Chryseobacterium, Sphingomonas, Pseudomonas, Thermoanerobacter, Methanothermobacter, Thermococcus, Petrotoga, Halomonas*, *Kosmotoga*, *Thermovirga*, *Archaeoglobus*, and *Caminicella* (Head, 2017; Kaster et al., 2009; Wentzel et al., 2013). Notably, despite being considered part of the indigenous community, *Thermovirga* was found in our samples exclusively in the samples after the water breakthrough, in the separators and in the injection water. This might suggest that this taxon might be oil processing contaminants rather than indigenous to the reservoir, or else, was present in low abundance within the reservoir and proliferated in the platform separators.

The predominant species in most samples were the strictly anaerobic bacterium *Thermoanaerobacter* and the archaeon *Thermococcus*. The MAGs taxonomic profiling based on the GTDB database (Fig. 7C) attributed the genus *Thermoanaerobacter* to two different species. *Thermoanaerobacter ethanolicus* (Wiegel and Ljungdahl, 1981), a non-spore-forming bacterium in the samples A18 before water breakthrough and the spore-former *Thermoanaerobacter pseudoethanolicus* (Onyenwoke et al., 2007), present in the wells A12 and A18 after water breakthrough. However, FastANI analysis (Jain et al., 2018) of the *Thermoanaerobacter* genomes suggests that all strains belong to the same species, as their pairwise ANI values exceed 95%, the species threshold defined by Jain et al. (2018). The observed discrepancy in GTDB classification might stem from differences in genome completeness, with marker genes in the more complete *Thermoaerobacter ethanolicus* genome influencing its placement. If the *Thermoanaerobacter pseudethanolicus* genome from A18 were more complete, it might also be classified as *Thermoanerobacter ethanolicus*. Alternatively, the missing genomic regions may genuinely distinguish *Thermoanaerobacter pseudethanolicus*, supporting the GTDB assignment.

*Thermoanaerobacter* survives within a temperature range of 35 to 80°C, with an optimum growth temperature of 65-70°C (Rainey and Stackebrandt, 1993). Similarly, *Thermococcus* is an obligate anaerobe and hyperthermophilic archaeon, with a known growth temperature range of 60°C to 105°C (Zhang et al., 2012).

### Effects of Water breakthrough on the community structure

The majority of taxa identified in the oil reservoir were also detected in the injection water. However, four genera - *Syntrophotalea*, *Desulfovibrio*, *Desulfofundulus*, and *Archaeoglobus* - were exclusive to the water injection and the water-injected wells. All four are anaerobic, thermophilic or thermotolerant and capable of sulphate-reduction. Their presence in post-injection oils can be attributed to their thermal resilience, as evidenced by their high optimal growth temperatures (e.g., *Archaeoglobus* at 83°C; Klenk et al., 1997). Among them, *Syntrophotalea* plays a significant role in degrading organic compounds such as butanol and ethanol (Sun et al., 2019). Additionally, *Syntrophotalea* can form syntrophic associations with methanogens, contributing to the degradation of crude oil, aromatic compounds, and alkanes (Gray et al., 2011).

We initially hypothesized that water breakthrough would significantly alter microbial distributions, leading to an overprinting of indigenous communities by injection-associated microbes. By contrast, reservoir engineers generally assume minimal mixing between injected water and oil, with water simply pushing oil ahead. However, while platform-associated microbial contamination was detected, extensive overprinting was absent. This is likely due to the short time since the water breakthrough (one month) and the inherent challenges faced by allochthonous microbes in adapting to reservoir conditions. Nonetheless, microbial diffusion within the reservoir appeared surprisingly rapid from a geological perspective.

Our metagenomic data closely mirrored the 16S rRNA gene data, revealing that the water separator and injection water shared similar viral genomic patterns and clustering. Given the high mutation rates of viruses compared to microorganisms, driven by their smaller genome sizes and rapid replication cycles (Sanjuán and Domingo-Calap, 2016), it is highly unlikely that identical viral strains would be found in isolated environments.

Microaerophilic respiration genes detected in the data do not imply active oxygen respiration but rather that the associated microorganisms can tolerate oxygen. These genes were scarce in pre-breakthrough samples but became more prevalent in post-breakthrough and injection water samples. This supports the notion that the taxa detected before the water breakthrough represent the indigenous microbial community of the oil reservoir.

### Methodological insights and future directions

A major difference between the 16S rRNA results and the metagenome results is a major underrepresentation of the archaeal taxa in the 16S rRNA data set. The genus *Thermococcus* in Fig. 2 seems less abundant than *Thermoanaerobacter,* whereas the metagenomic data in Fig. 7B show the pattern reversed. This discrepancy highlights the known biases in 16S rRNA gene sequencing, where primer design often favours bacterial sequences, leading to poor amplification of archaeal 16S rRNA genes (Teske and Sørensen, 2008). By contrast, the *rps3* gene sequence obtained through the metagenomes provides a more accurate picture of microbial diversity. Our findings emphasize the need to complement 16S rRNA studies with metagenomic approaches if the study aims to quantitatively identify microbial community structure.

Accurately characterizing oil reservoir microbial communities requires analysing crude oil before any separation processes occur (Alibrandi et al., 2025, in review). Our data indicate that the taxonomic composition of injection water closely mirrors that of both the water and oil separators (Fig. 4), suggesting that contamination may derive from biofilms on pipe surfaces and within the separators. This observation implies that either allochthonous microbes adapt to reservoir conditions or that rare reservoir indigenous microbes preferentially proliferate in the platform separator and associated piping. Moreover, despite the injection water containing seawater, seawater-associated taxa were scarce, which we attribute to the high-temperature treatment and the addition of biocides during injection that likely eradicate most seawater-derived organisms. Overall, our data reveal that the water injection, reservoir, and oil platform infrastructure communities are intricately intertwined and shape the final microbial composition of all samples as evidenced by the relative 16S rRNA gene abundance at the genus level (Fig. 5) and IndVal results (Fig. 6). It is not surprising that early studies of microbial communities associated with oil reservoirs often questioned the origin of the observed taxa.

Our findings contribute to the broader understanding of the microbial ecology in high-temperature oil reservoirs and highlight the remarkable resilience of certain genera facing extreme thermal conditions. While oil reservoir microbiology has historically been overlooked by the oil industry, this study underscores its significance as a valuable parallel tool for oil field monitoring. Future research endeavours should delve deeper into the long-term impacts of flooding throughout the entire lifespan of reservoirs from discovery to decommissioning. These insights into microbial diversity and community structure in a deeply buried, polyextreme environment have important implications for reservoir management strategies.

## Acknowledgements

We would like to thank Aker BP for providing funding and samples for this project, and our lab technicians at GeoForschungsZentrum Potsdam, Anke Sabowrowski, Oliver Burckhardt and Axel Kitte for their support. I would also like to thank Lucas Horstmann for the immense help with R scripts.

## Ethics statement

This study did not involve any trials on humans or animals

## Data availability statement

The sequencing data is available via the ENA (European Nucleotide Archive) under the project accession PRJEB81118.

