## supplementary figures for "Microbial Diversity and Community Shifts in a Petroleum Reservoir under Production: Effects of Water Breakthrough and Anthropogenic Alterations"

(4) Aker BP, Norway.

**Content:** supplementary methods, supplementary figures and tables, supplementary references.

**Supplementary methods**

*Phylogenetic tree based on 16S rRNA genes*

For phylogenetic analysis of 16S rRNA genes, a Maximum Likelihood (ML) tree was constructed using RAxML to examine the partial sequences of the V4 hypervariable region of 16S rRNA genes. Taxonomic assignments focused on groups relevant to oil reservoir environments, specifically targeting sulfate-reducing bacteria and extremophilic lineages within Bacteria, and methanogens and extremophilic groups within Archaea.

*Viral metagenome method*

Putative viral scaffolds in the metagenomes were identified using three different tools: VIBRANT v. 1.2.1(Kieft et al., 2020); VirSorter2 v. 2.2.4 (Guo et al., 2021) in sensitive mode; and DeepVirFinder v.1.0 (Ren et al., 2020) with a threshold of 0.7. Hosts contaminations were removed and completeness was calculated using CheckV v. 1.0.1 (Nayfach et al., 2021). Only scaffolds, predicted as viral by all three tools, with predicted completeness ≥ 25 % and no warnings by CheckV v. 1.0.1 were used for further analyses. The putative viral scaffolds were mapped with Bowtie2 v. 2.4.1 (Langmead and Salzberg, 2012) and samtools v. 1.13 (Li, 2009) against all quality-controlled raw reads. We used the mappings to calculate and compare single nucleotide polymorphism (SNPs) and average nucleotide identity (ANI) with InStrain v. 1.8.1 (Olm et al., 2021) between the viral strains.

**Supplementary Figures**

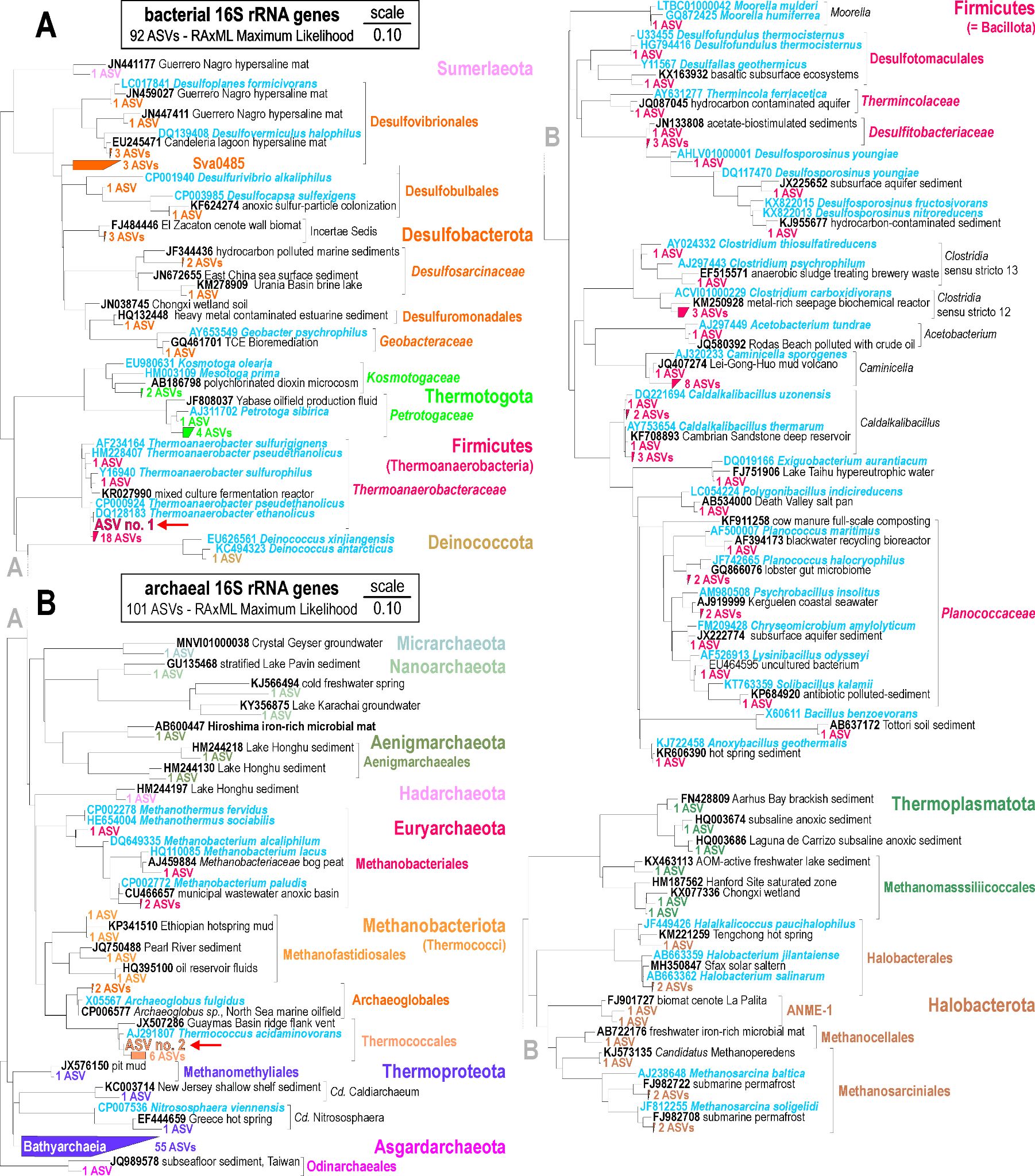
**Supplementary figure S1: Phylogenetic analyses of 16S rRNA genes.** RaxML Maximum Likelihood phylogenetic tree of partial 16S rRNA genes (V4 hypervariable region) taxonomically assigned to (**A**) sulfate reducers and extremophiles among Bacteria and (**B**) methanogens and extremophiles among Archaea.

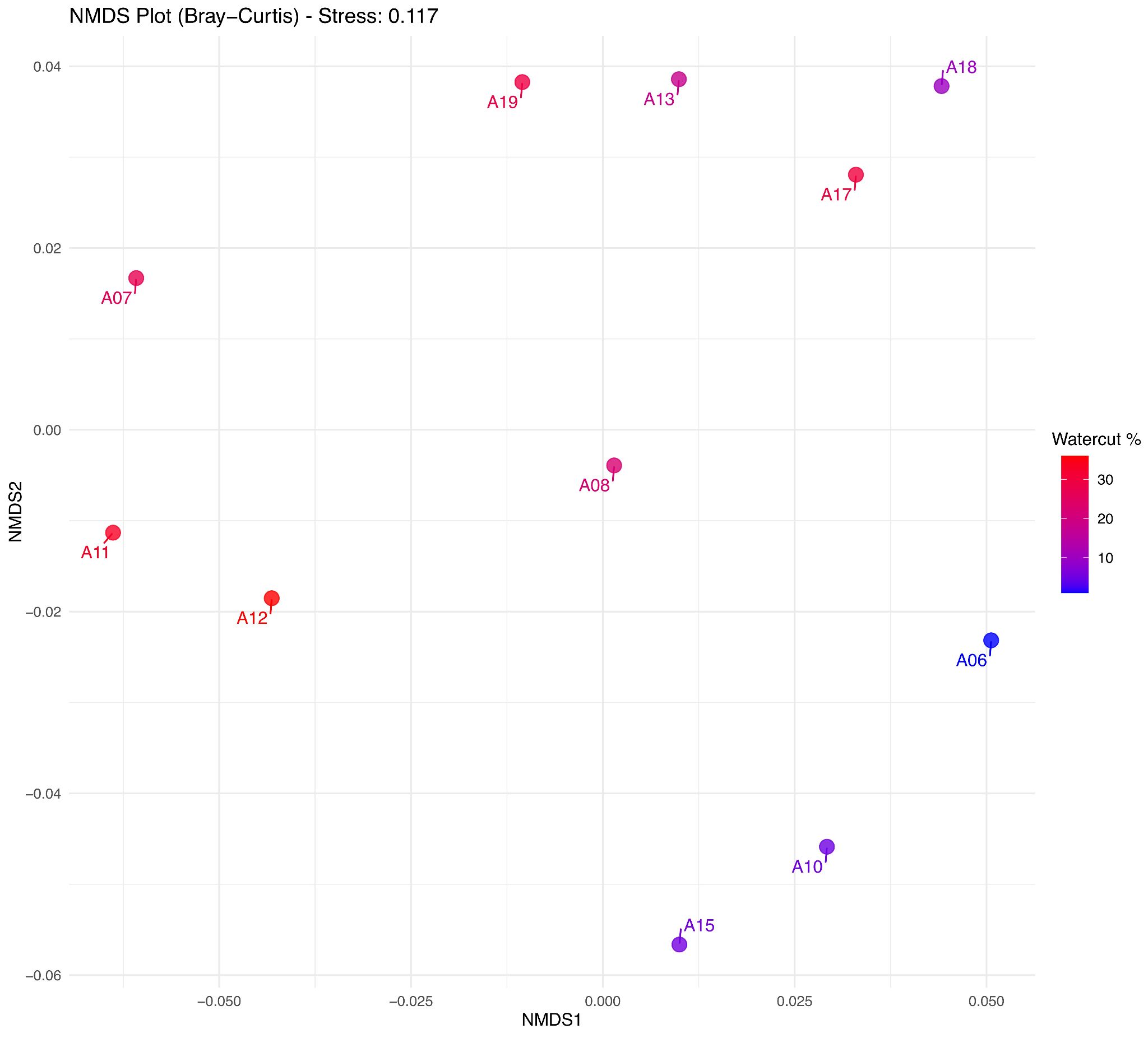

**Supplementary Figure S2: Non-metric multidimensional scaling (NMDS) plot based on Bray-Curtis dissimilarity, illustrating microbial community composition across samples.** Each point represents a sample, colored according to its Watercut value (blue = lower Watercut, red = higher Watercut). The proximity of points reflects the similarity of microbial communities, with closer points indicating more similar compositions. The stress value (0.117) indicates the goodness of fit for the ordination. A gradient in color distribution suggests a potential relationship between Watercut and microbial community structure.

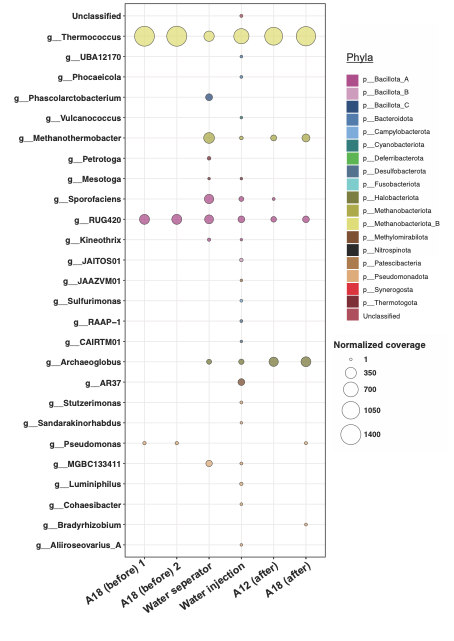
**Supplementary figure S3: Normalized coverage of extended rpS3 gene sequences summarized for the assigned genera.** Bubbles are coloured according to phyla.

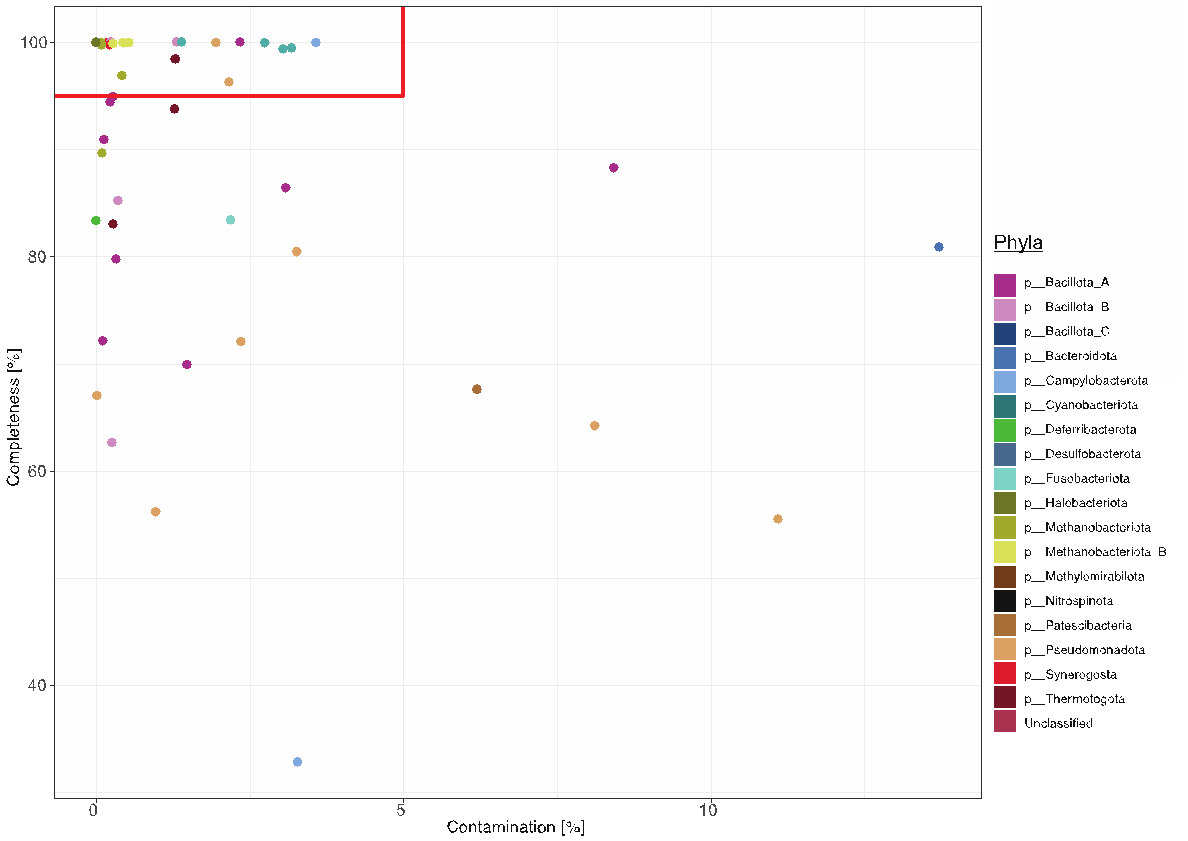

**Supplementary Figure S4: Completeness and contamination of metagenome assembled genomes (MAGs).** Red rectangular marks thresholds for high quality MAGs. Color indicates phyla of MAG.

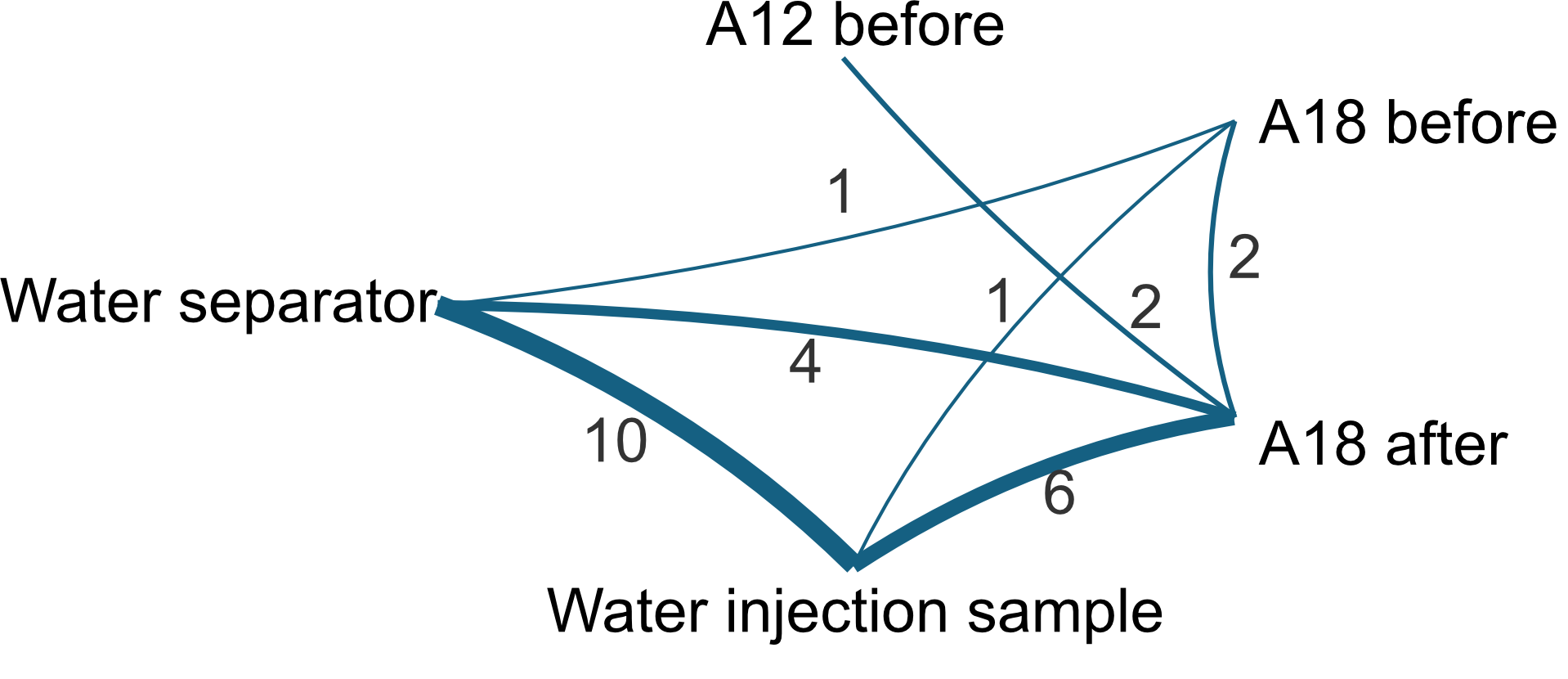

**Supplementary figure S5: Edvard Grieg viral strain clustering.** Line thickness corresponds to shared viral strain clusters (also displayed as numbers on the lines).

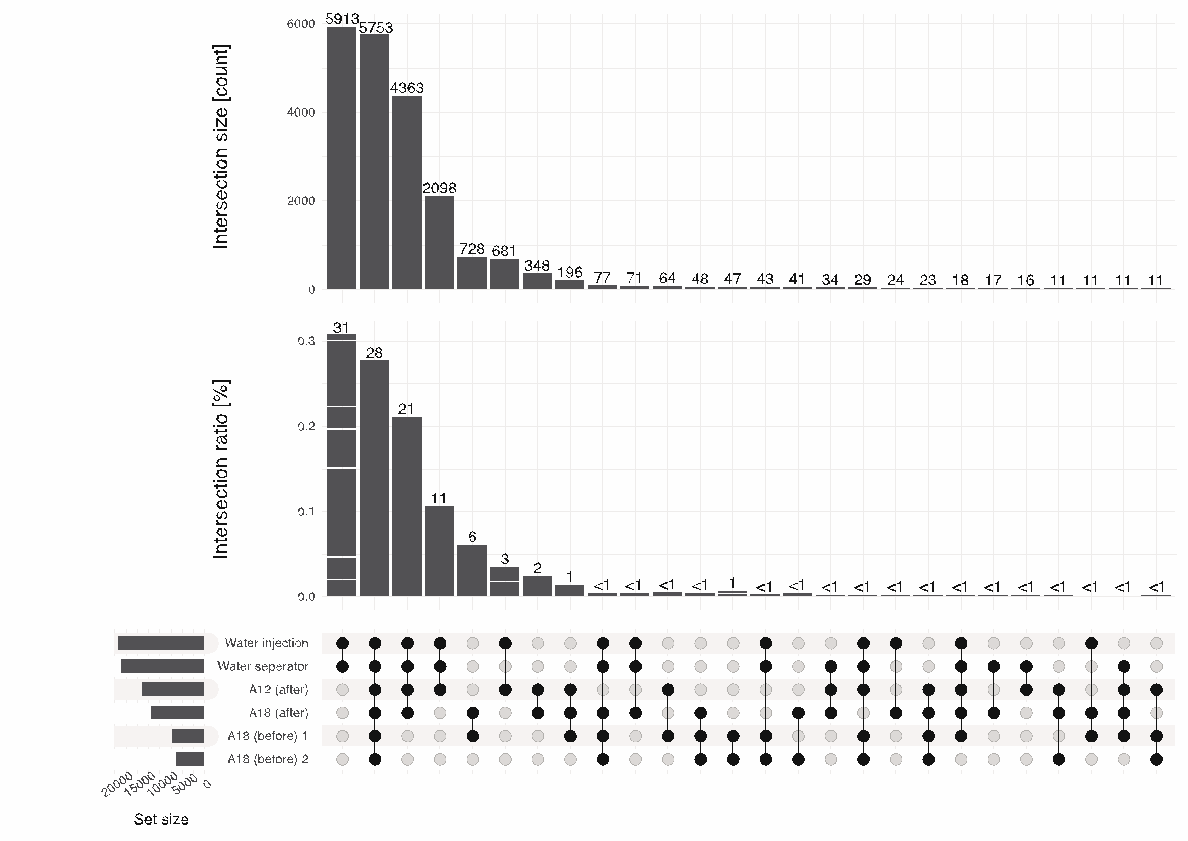

**Supplementary Figure S Intersection size and percent intersection ratio (proportion of shared cluster between the respective intersection) of the non-singleton gene cluster (total 20,769) between the six metagenomes.** All sets with more than 10 shared clusters are shown.

**Supplementary table S1:** Permanova analysis results comparing the differences between the wells. The (*) indicates the samples that show no differences.

**Supplementary Table S2**: Permanova analysis results comparing the differences between water injection, water separator oil separator and the oil samples from wells A12 and A18 before and after fluid breakthrough. The (*) indicates the samples that show no differences.

**Supplementary Table S3:** Metadata of the assembly of metagenomic reads into scaffolds for the six samples whose libraries were successfully sequenced.

**Supplementary Table S4:** Metadata on the *de novo* assembly of the 52 metagenome-assembled genomes (MAGs) obtained from the six metagenomic assemblies.

**Supplementary Table S5:** Coverage of extended *rpS3 gene* sequences in eleven metagenomic samples and GTDB taxonomy (Parks et al. 2022) of *rp3 gene* sequences. Coverage is only given for mappings with a minimum breadth of 95%.

**Supplementary Table S6:** Coverage normalized by base pair count of the forward reads of protein cluster representatives in the six metagenomic samples.

**Supplementary Table S7:** number of genes of interest hits from the BlastP data table.

**Supplementary Table S8:** Permanova analysis results comparing the difference between the ORFs of oils before and after injection, water separator and water injection.

***References***

Bushnell, B.: BBMap: a fast, accurate, splice-aware aligner, 2014.

Cantalapiedra, C. P., Hernández-Plaza, A., Letunic, I., Bork, P., and Huerta-Cepas, J.: eggNOG-mapper v2: functional annotation, orthology assignments, and domain prediction at the metagenomic scale, Mol. Biol. Evol., 38, 5825–5829, 2021.

Graham, E., Heidelberg, J., and Tully, B.: Potential for primary productivity in a globally-distributed bacterial phototroph, ISME J., 12, 1861–1866, 2018.

Guo, J., Bolduc, B., Zayed, A. A., Varsani, A., Dominguez-Huerta, G., Delmont, T. O., Pratama, A. A., Gazitúa, M. C., Vik, D., and Sullivan, M. B.: VirSorter2: a multi-classifier, expert-guided approach to detect diverse DNA and RNA viruses, Microbiome, 9, 1–13, 2021.

Hyatt, D., Chen, G.-L., LoCascio, P. F., Land, M. L., Larimer, F. W., and Hauser, L. J.: Prodigal: prokaryotic gene recognition and translation initiation site identification, BMC Bioinformatics, 11, 1–11, 2010.

Jiang, H., Lei, R., Ding, S.-W., and Zhu, S.: Skewer: a fast and accurate adapter trimmer for next-generation sequencing paired-end reads, BMC Bioinformatics, 15, 1–12, 2014.

[Joshi, N. and Fass, J.: Sickle: A sliding-window, adaptive, quality-based trimming tool for FastQ files (Version 1.33)[Software], 2011.](https://www.zotero.org/google-docs/?0YwWLA)

Kang, D. D., Li, F., Kirton, E., Thomas, A., Egan, R., An, H., and Wang, Z.: MetaBAT 2: an adaptive binning algorithm for robust and efficient genome reconstruction from metagenome assemblies, PeerJ, 7, e7359, 2019.

Kieft, K., Zhou, Z., and Anantharaman, K.: VIBRANT: automated recovery, annotation and curation of microbial viruses, and evaluation of viral community function from genomic sequences, Microbiome, 8, 1–23, 2020.

Kieser, S., Brown, J., Zdobnov, E. M., Trajkovski, M., and McCue, L. A.: ATLAS: a Snakemake workflow for assembly, annotation, and genomic binning of metagenome sequence data, BMC Bioinformatics, 21, 1–8, 2020.

Langmead, B. and Salzberg, S. L.: Fast gapped-read alignment with Bowtie 2, Nat. Methods, 9, 357–359, 2012.

Letunic, I. and Bork, P.: Interactive Tree of Life (iTOL) v6: recent updates to the phylogenetic tree display and annotation tool, Nucleic Acids Res., gkae268, 2024.

Li, H.: The sequence alignment/map (SAM) format and SAMtools 1000 genome project data processing subgroup, Bioinformatics, 25, 1, 2009.

Nayfach, S., Camargo, A. P., Schulz, F., Eloe-Fadrosh, E., Roux, S., and Kyrpides, N. C.: CheckV assesses the quality and completeness of metagenome-assembled viral genomes, Nat. Biotechnol., 39, 578–585, 2021.

Nurk, S., Meleshko, D., Korobeynikov, A., and Pevzner, P. A.: metaSPAdes: a new versatile metagenomic assembler, Genome Res., 27, 824–834, 2017.

Olm, M. R., Crits-Christoph, A., Bouma-Gregson, K., Firek, B. A., Morowitz, M. J., and Banfield, J. F.: inStrain profiles population microdiversity from metagenomic data and sensitively detects shared microbial strains, Nat. Biotechnol., 39, 727–736, 2021.

Parks, D. H., Imelfort, M., Skennerton, C. T., Hugenholtz, P., and Tyson, G. W.: CheckM: assessing the quality of microbial genomes recovered from isolates, single cells, and metagenomes, Genome Res., 25, 1043–1055, 2015.

Parks, D. H., Chuvochina, M., Rinke, C., Mussig, A. J., Chaumeil, P.-A., and Hugenholtz, P.: GTDB: an ongoing census of bacterial and archaeal diversity through a phylogenetically consistent, rank normalized and complete genome-based taxonomy, Nucleic Acids Res., 50, D785–D794, 2022.

Ren, J., Song, K., Deng, C., Ahlgren, N. A., Fuhrman, J. A., Li, Y., Xie, X., Poplin, R., and Sun, F.: Identifying viruses from metagenomic data using deep learning, Quant. Biol., 8, 64–77, 2020.

Sieber, C. M., Probst, A. J., Sharrar, A., Thomas, B. C., Hess, M., Tringe, S. G., and Banfield, J. F.: Recovery of genomes from metagenomes via a dereplication, aggregation and scoring strategy, Nat. Microbiol., 3, 836–843, 2018.

Wu, Y.-W., Simmons, B. A., and Singer, S. W.: MaxBin 2.0: an automated binning algorithm to recover genomes from multiple metagenomic datasets, Bioinformatics, 32, 605–607, 2016.
